# How stra(i)nge are your controls? A comparative analysis of metabolic phenotypes in commonly used C57 substrains

**DOI:** 10.1101/2023.03.03.530990

**Authors:** Annesha Sil, Marina Souza Matos, Mirela Delibegovic, Bettina Platt

## Abstract

In recent years, the use of insufficiently characterised control subjects has been a contributing factor to increasing irreproducibility in different areas of biomedical research including neuroscience and metabolism. There is now a growing awareness of phenotypic differences between the metabolic profiles of C57BL/6 substrains which are commonly used as control animals.

We here investigated baseline metabolic characteristics such as glucose regulation, fasted serum insulin levels and hepatic insulin signalling in five different C57BL/6 sub-strains (N, J, JOla, JRcc) of both sexes, obtained from two commercial vendors Charles River Laboratories (Crl) and Envigo (Env).

Our results indicated systematic and tissue-specific differences between substrains, modulated by both vendor and sex in all parameters investigated, not necessarily mediated by the presence of the *Nnt* mutation. Not only were there differences between 6J and 6N as expected, all three 6J sub-strains exhibited different profiles, even from the same breeder. Two distinct metabolic profiles were identified, one in which low insulin levels resulted in impaired glucose clearance (6JCrl; both sexes) and the other, where sustained elevations in fasted basal insulin levels led to glucose intolerance (male 6JRccEnv). Further, 6JRccEnv displayed sex differences in both glucose clearance and hepatic insulin signalling markers. In comparison, the two 6N substrains of either sex, irrespective of vendor, did not exhibit considerable differences, with 6NCrl animals presenting a good choice as a healthy baseline ‘control’ for many types of experiments.

Overall, our data emphasise the importance of selecting and characterising control subjects regarding background, sex, and supplier to ensure proper experimental outcomes in biomedical research.

## INTRODUCTION

In recent years, biomedical research has found itself at the centre of a reproducibility crisis [1]. Among the many issues identified, one crucial factor is the use of insufficiently characterised controls, especially in studies involving the use of animals, which can distort experimental outcomes and conclusions drawn. Concurrently, there has been increasing awareness of the differences between inbred mouse substrains used in different areas of biomedical research, including neuroscience and metabolic research [2]. Here, ‘inbred’ strains are defined as any strain where brother-sister mating has occurred for at least 20 consecutive generations, whereas ‘substrain’ is a genetically distinct branch that has significantly deviated from the original founding strain. One of the most commonly used inbred mouse strains in biomedical research are the C57BL/6 lines, which can be purchased from multiple commercial vendors around the globe, including Charles River (Crl), Envigo (Env; Harlan Laboratories), The Jackson Laboratory (JAX), Taconic, Janvier, and Animals Resource Centre (Arc) [3]. The C57BL/6J (6J) and C57Bl/6N (6N) substrains are usually the primary choice for many types of biomedical studies due to ease of breeding and background stability, which also make these a good choice for creation of genetically altered animals [4]. Published literature using these ‘wild-type’ strains often do not take into consideration the genetic make-up and phenotypic differences already reported [5–7] and are often plagued by insufficient information or nomenclature; offering only “C57” or “C57BL6” labels, without further details.

Several single nucleotide polymorphisms (SNPs) have been identified between 6J and 6N mice [8]. Regulatory and coding regions containing SNPs can severely change gene expression and function [6]. In 2005, a spontaneous in-frame five-exon deletion in the *Nnt* gene was identified in 6J mice [5]. The affected nicotinamide (NAD) nucleotide transhydrogenase (NNT) protein is a mitochondrial enzyme responsible for the reduction of NADP^+^ to NADPH, necessary for ATP synthesis. NAPDH detoxifies reactive oxygen species (ROS) in the mitochondria, essential for H_2_O_2_ clearance and redox homeostasis. Consequently, the *Nnt* mutation has been linked to metabolic phenotypes observed in 6J mice, including impaired insulin secretion and glucose intolerance, correlated with major redox imbalances [5,9,10].

At present, conflicting data exist regarding metabolic differences between 6J and 6N mice including their glucose tolerance and body weight [4]. Previous studies demonstrated that 6J and 6N animals develop glucose intolerance when on high-fat diet (HFD), while 6J mice showed higher glucose levels [11] and lower insulin secretion than 6N mice [12]. However, other studies have reported no major differences in glucose tolerance or insulin secretion between 6J and 6N [4,13,14]. Hence, *Nnt* is unlikely to be the only genetic factor contributing to metabolic phenotypes.

There is also evidence concerning phenotypic differences between the 6J substrains from different suppliers. SNP analysis showed that 6JCrl, 6JArc, and 6J (JAX) were genetically identical, but those from other suppliers displayed substrain-specific polymorphisms other than Nnt that could alter behaviour [15]. For example, 6JOlaHsd (Envigo) mice have a loss-of-function deletion in the alpha-synuclein (*Scna*) and multimerin 1 (*Mmrn1*) genes. While the deletion of *Scna* does not seem to impact learning, it nonetheless has important implications for Parkinson’s Disease (PD) research [16,17]. Moreover, the *Mmrn1* mutation promotes impaired platelet adhesion and thrombus formation [18], crucial for e.g. wound healing and thrombosis research. Remarkably, all 6N substrains show similar genetic profiles independent of the supplier [19], but display a nonsense mutation in the *Crb1* gene that promotes retinal degeneration 8 (*rd8* mutation), which complicates not only ophthalmological studies but also behavioural testing [4].

So far, hepatic insulin signalling central for systemic glucose homeostasis has not been characterised adequately in C57 substrains. Altered insulin signalling can ultimately lead to insulin resistance, hyperglycaemia, and type 2 diabetes (T2D) [26]. Insulin receptor (IR) activation triggers phosphorylation of protein kinase B (AKT) via phosphoinositide-3-phosphate kinase (PI3K). AKT can regulate multiple pathways to regulate glucose and lipid homeostasis, e.g. glycogen synthesis and inhibition of the glycogen synthase kinase 3β (GSK3β). Additionally, activated AKT can lead to an increase in mammalian target of rapamycin complex (mTOR), further facilitating the phosphorylation of p70 ribosomal S6 kinase 1 (S6K1) and activation of the ribosomal protein S6 (rpS6), which promotes protein synthesis. rpS6 activation also plays an important role in insulin receptor substrate 1 (IRS-1) inhibition and insulin resistance [27,28].

In this study, we demonstrate systemic and tissue-specific differences in glucose regulation, baseline fasted insulin levels and hepatic insulin signalling between five C57BL/6 substrains of both sexes, obtained from different commercial vendors. Our data imply that multiple factors affect the metabolic phenotypes observed and stress the importance of selecting and characterising appropriate controls. We highlight the requirement to correctly and comprehensively describe background, sex, and supplier of animals used, as these can be confounders for several fields of biomedical research.

## MATERIALS AND METHODS

This work was conducted according to the ARRIVE guidelines 2.0 for reporting animal research [20] and the EQIPD framework for the internal validity in the design, conduct, and analysis of preclinical biomedical experiments involving laboratory animals [21].

### Animals

4-month-old male and female C57BL/6 mice belonging to two different substrains (broadly 6J or 6N) were obtained from two different suppliers: Charles River (Tranent, UK) and Envigo (Netherlands). C57BL/6JCrl (6JCrl) and C57BL/6NCrl (6NCrl) were supplied by Charles River while two different 6J substrains - C57BL/6JOlaHsd (6JOlaEnv), C57Bl/6JRccHsd (6JRccEnv), and a 6N substrain C57BL/6NHsd (6NEnv) were supplied by Envigo (Table 1). A total of 60 animals (Females: n=4 for 6JCrl, 6NCrl, 6NEnv, n=4 for 6JOlaEnv, n=10 for 6JRccEnv; males: n=6 for 6JCrl, 6NCrl, 6JOlaEnv, 6NEnv, n=10 for 6JRccEnv) were utilised for experiments. All animals were delivered to the Medical Research Facility (University of Aberdeen) at least two weeks prior to experimentation for habituation. Animals were group-housed in stock cages (Tecniplast, Italy), and maintained on a 12-h day-night cycle (lights on 7am, simulated dusk/dawn 30 min) in temperature- (20–22°C) and humidity- (60–65%) controlled holding facilities with *ad libitum* access to standard chow (Special Diet Services, Witham, UK) water and enrichment exactly as described previously [22]. Animals were tail-handled multiple times a week to reduce anxiety prior to testing. Glucose tolerance testing (GTT) took place on weekdays during the light period (09:00–16:00) in a separate testing room. Genotyping for *Nnt* knockout or *Nnt* wildtype mutation in all substrains was carried out by Transnetyx USA and confirmed that as expected, only 6JCrl group had the *Nnt* knockout mutation.. All procedures were approved by local ethical review, a UK Home Office project license and complied with the EU directive 63/2010E and the UK Animal (Scientific Procedures) Act 1986.

**Table 1:**
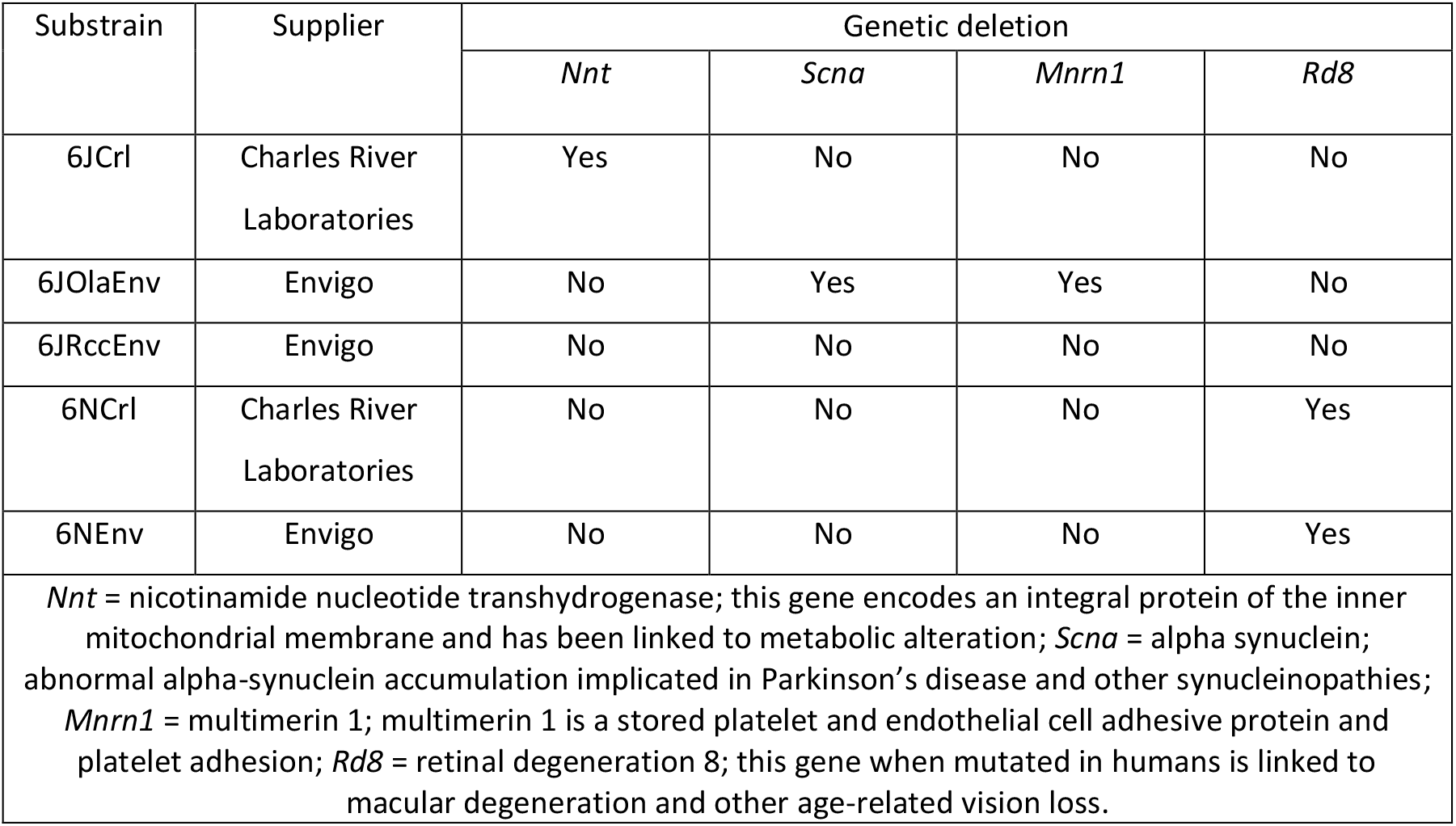
List of different substrains utilised with some important known genetic deletions

### Glucose Tolerance Test (GTT)

Mice were fasted for 5 hours before GTT between 08:00-13:00 on weekdays with *ad libitum* access to water maintained as described previously [23]. Fasting blood glucose was measured by tail puncture (time 0) prior to intraperitoneal injection of glucose solution (2mg/g body weight). Blood glucose was assessed at 15-, 30-, 60-, and 90-minutes post-injection. AlphaTRAKII veterinary glucometer (Berkshire, UK) was used to determine blood glucose levels. The glucometers were calibrated in accordance with instructions prior to testing. One female 6JOlaEnv and one female 6JRccEnv mouse did not complete the experiment as they had low glucose levels and were excluded from the analysis. Total glycaemic excursion as measured by area under curve (AUC) as well as blood glucose concentration (mmol/L) at time intervals listed above were plotted and analysed.

### Tissue extraction and sample preparation

After a 5-hour fast, animals were humanly culled using cervical dislocation. Extracted livers were snap-frozen in liquid nitrogen and stored in –80°C until use. For serum insulin measurements, blood derived from the trunk was collected into separator micro-tubes (BD Microtainer, Canada) and allowed to clot at room temperature for approximately 15 minutes. Tubes were centrifuged at 7,500 rpm for 15 min at 4°C. Serum aliquots were moved to a new tube and stored in -80°C freezer until use. For western blotting experiments, liver tissues were homogenised in RIPA buffer (10 mM Tris-HCl, 150 mM NaCl, 0.1% SDS, 1% Triton, 1% sodium deoxycholate, 5 mM EDTA: pH = 7.4) supplemented with protease inhibitors (Roche) and PhosStop tablets (Roche). The homogenates were centrifuged at 12,000 rpm for 20 min at 4°C; the supernatants constituted the soluble fraction which was used in immunoblotting experiments.

### Serum insulin ELISA

Serum insulin concentrations were determined using ultra-sensitive mouse insulin ELISA kit (CrystalChem, UK), accordingly to the manufacturer’s protocol [23].

### Immunoblotting

Western blots were performed as described previously [22,24]. In short, protein concentration was adjusted (3 μg/ml) following bicinchoninic acid assay measurement (BCA, SigmaAldrich, Poole, UK) and dilution in lysis buffer. Cell lysates were prepared for electrophoresis (RIPA buffer, 4X lithium dodecyl sulphate (LDS, Thermo Fisher Scientific, UK), and 15mM dithiothreitol (DTT)) and heated for 10 min at 70°C, followed by separation for 45 min at 200 V in MOPS buffer in pre-cast 4 – 12% gradient Bis-Tris precast gels (Nupage, Thermo Fisher Scientific). This was followed by transferring onto nitrocellulose membranes (0.45-μm pore size, Invitrogen, UK) for 1h at 25V, at room temperature. Following this, membranes were blocked with 5% BSA-TBST (0.05% Tween, 50mMTrizma base, 150mMNaCl) for phosphorylated (p-) proteins or 5% milk-TBST for total (t-) proteins, for 1 h. Following TBST washes, membranes were incubated with primary antibodies prepared in 5% BSA, 0.05% sodium azide, and TBST (p-AKT (Ser473): Cell Signalling #4060, 1:1000, t-AKT: Cell Signalling #4691, 1:1000, p-rpS6 (Ser235/237): Cell Signalling #4858 1:1000, t-rpS6: Cell Signalling #2217, 1:1000, p-GSK3β (Ser9): Cell Signalling #9323, 1:500, t-GSK3β: Cell Signalling #9315, 1:1000) overnight at 4°C. Next, they were incubated in their respective secondary antibodies for 1 h at room temperature (goat anti-rabbit, IgG, HRP conjugated; Merck Millipore (1: 5000) in 5% milk-TBST) and visualised using enhanced chemiluminescent substrate (ECL; 0.015% hydrogen peroxide (H_2_O_2_), 30 μM coumeric acid in 1.25 mM luminol). An iBright™ FL1000 Imaging System camera (Invitrogen™, ThermoFisher Scientific) was utilized to capture immunoreactivity at 16bit for analysis. For probing of t-protein after visualisation of p-proteins, membranes were washed with a mild stripping buffer (59.52 mM glycine, 81 μM SDS, 1% Tween 20; pH 2.2). Membranes were also processed for total protein using both Ponceau-S stain as described previously [22] to control for loading. Densitometric analysis was performed as area under the curve (AUC) using ImageJ software (Ver. 1.53, NIH, USA) and normalised as per Ponceau staining (both p- and t-). Additionally, the ratio of p- to t-protein was calculated for all pathways investigated. In accordance with our previous publications, fold increase of the protein of interest was then calculated relative to 6JCrl as they are very commonly used in biomedical research as controls [22].

### Statistical analysis

Statistical analysis for GTTs, insulin ELISA and western blots was performed using one-way Analysis of Variance (or Welch’s ANOVA when standard deviation (SDs) between groups was different) followed by Tukey’s post-hoc tests in females and males separately (Prism, Version 8.4, GraphPad USA). Body weight comparisons were carried out using a regular two-way ANOVA with substrain and sex as factors. As it was the intention of the authors to also examine potential sex differences within substrains, if sex-dependent phenotypes were observed in any of the parameters examined, pre-planned comparison t-tests between male and female animals of the same substrain were performed. Data are presented as scatter plots with means ± SD. For all analyses, alpha was set to 5% and p < 0.09 considered as ‘marginally significant’ as per recent recommendations [25]. It is acknowledged that in the female cohort, low n’s maybe a contributing factor for the observance of marginal significances rather than outright significant differences.

### Correlational analysis

Correlational analysis using GraphPad Prism was carried out between AUC during GTT, serum insulin levels and molecular markers of interest in the different substrains in male animals as this provided coherent groups with the greatest power (greater n’s compared to females). All raw data underwent Z-transformation within each group (Microsoft Excel 365) followed by Pearson’s correlation analyses after confirmation of Gaussian distribution of datasets. Correlations are displayed as heat plot matrices with red indicating negative correlations and blue indicating positive ones. Values of individual correlational coefficients and their p-values up to 0.09 (dark green to light green for p-values between 0.001-0.1), are represented in the Supplementary Material. For all comparisons, significance was set at p < 0.05.

## RESULTS

### Differences in glucose tolerance between C57BL/6 substrains in both sexes

Basal glucose levels and total glycaemic excursion via GTTs were assessed in both male and female mice from different C57BL/6 sub-strains. As expected, body weights of the males overall were higher than females (F (1,48) = 213.4, p<0.0001), and no differences in body weight between substrains were observed (F (4,48) =1.615, p=0.18) (Fig. 1A).

**Fig. 1:**
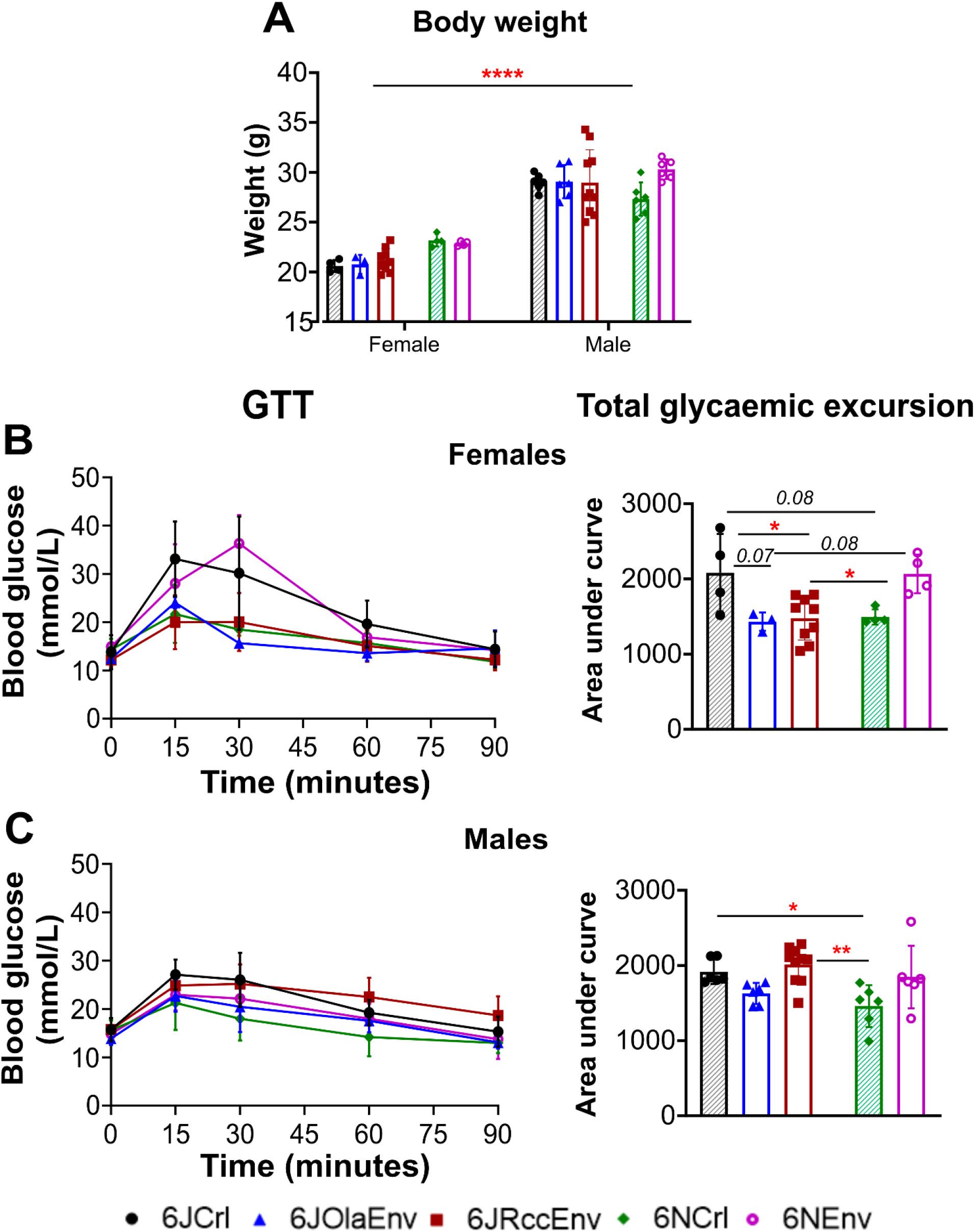
Body weight and GTT profiles of C57 substrains of both sexes: Body weight (**A**) of male and female animals belonging to five C57 substrains i.e., 6JCrl (black with shaded pattern), 6JOlaEnv (blue), 6JRccEnv (maroon), 6NCrl (green with shaded pattern), 6NEnv (purple) where male animals had higher bodyweight overall compared to females with no substrain differences (regular 2-way ANOVA with substrain and sex as factors). Blood glucose levels (mmol/L) measured every 15 minutes (left) and total glycaemic excursion measured as total area under curve (right) during the 90-minutes of the GTT for females (B) and male (C) C57 substrains. Differences in glycaemic excursion between substrains within each sex interrogated using one-way ANOVA with Tukey’s post-hoc tests such that 6JCrl, 6NEnv (both sexes) and 6JRccEnv (males) had highest total glycaemic excursion. All data sets visualised as scatter plots with mean ± SD with significances for displayed on graph as: **=p<0.01, *=p<0.05 (Tukey’s post-hoc test), p<0.09 in italics for marginally significant differences. Key for each substrain displayed at the bottom of the figure with Crl (Charles River Laboratories) animals of J and N substrain having shaded pattern inside to distinguish from Env (Envigo).

Importantly, significant differences in glucose tolerance were noted in both sexes (Females: F (4,19) = 5.34 p=0.004); Males: (F (4,29) = 5.10, p=0.003). In females, 6JCrl were found to be either significantly or marginally significantly *less* glucose tolerant than most other substrains apart from 6NEnv (Fig. 1B) as displayed by higher area under curve during the GTT (6JCrl vs 6JOlaEnv: p=0.07; 6JCrl vs 6JRccEnv: p=0.02; 6JCrl vs 6NCrl: p=0.08). Meanwhile, 6NEnv showed higher total glycaemic excursion (indicating *less* glucose tolerance) compared to 6JRccEnv (p=0.02) and 6JOlaEnv (p=0.08).

In male animals (Fig 1C), it was the 6JRccEnv line which was *more* glucose intolerant compared to the 6JOlaEnv (p=0.05) and 6NCrl (p=0.002). Additionally, the 6JCrl mice had higher total glycaemic excursion vs. 6NCrl (p=0.04). A planned paired comparison t-test between male and female 6JRccEnv animals confirmed that male 6JRccEnv animals were much more glucose intolerant compared to the females (p=0.0004).

Further, the basal glucose levels after 5h of fasting were only different between female substrains (Fig. 2A; Welch’s ANOVA: F (4.000, 7.851) = 4.013, p=0.04) but not in males (Fig 2B; F (4,29) =1.68, p=0.14). For females, 6NEnv had higher basal fasted glucose levels compared to 6JRccEnv (p=0.03) and 6JOlaEnv (p=0.08, marginally significant).

**Fig. 2:**
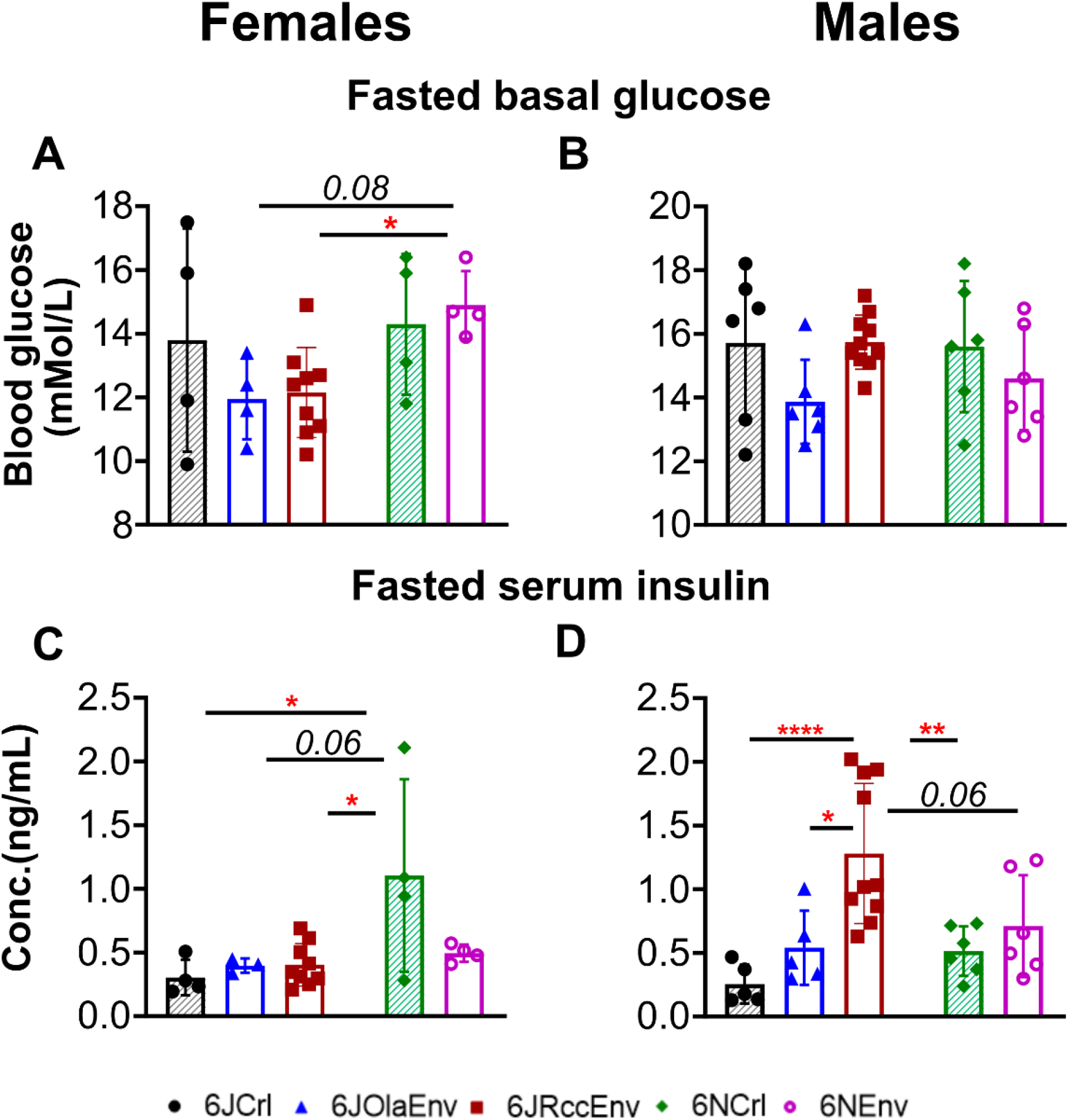
Basal fasted glucose and serum insulin levels in C57 substrains of both sexes: Basal glucose and serum insulin levels after a 5hr fast in female (**A, C** respectively) and male (**B**,**D** respectively) animals belonging to five C57 substrains i.e., 6JCrl (black with shaded pattern), 6JOlaEnv (blue), 6JRccEnv (maroon), 6NCrl (green with shaded pattern), 6NEnv (purple). Differences in glucose/insulin levels between substrains within each sex interrogated using one-way ANOVA with Tukey’s post-hoc tests such that there were no differences in fasted basal glucose levels between male substrains and 6NEnv female animals had higher fasted basal glucose compared to 6JRccEnv and 6JCrl; 6JCrl animals of both sexes had very low while 6JRccEnv males had highest basal fasted serum insulin levels. All data sets visualised as scatter plots with mean ± SD with significances for displayed on graph as: ****=p<0.0001, **=p<0.01, *=p<0.05 (Tukey’s post-hoc test), p<0.09 in italics for marginally significant differences. Key for each substrain displayed at the bottom of the figure with Crl (Charles River Laboratories) animals of J and N substrain having shaded pattern inside to distinguish from Env (Envigo).

### Strain and sex differences in fasting serum insulin levels

In female mice, 6NCrl female mice exhibited higher serum insulin levels (F(4,19)=4.071) compared to 6JRccEnv (p=0.01), 6JOlaEnv (p=0.06) and 6JCrl (p=0.01), but not 6NEnv (Fig. 2C). A different profile was observed in male animals, where 6JRccEnv displayed the highest serum insulin levels after 5 hours of fasting (F (4,40) = 8.099) compared to 6JOlaEnv (p=0.03), 6JCrl (p<0.0001), and 6NCrl (p=0.002) (Fig. 2D).

### Strain and sex differences in hepatic insulin signalling

Western blot analysis of the AKT pathway revealed differences between substrains only in male animals (Fig 3F; females are shown in Fig 3B) for p-AKT (F(4,9)=1.15, p=0.39), t-AKT (F(4,10)=1.690, p=0.22) and the ratio of p-AKT/ t-AKT (p/t-AKT; F(4,10)=1.909, p=0.18).

**Fig. 3:**
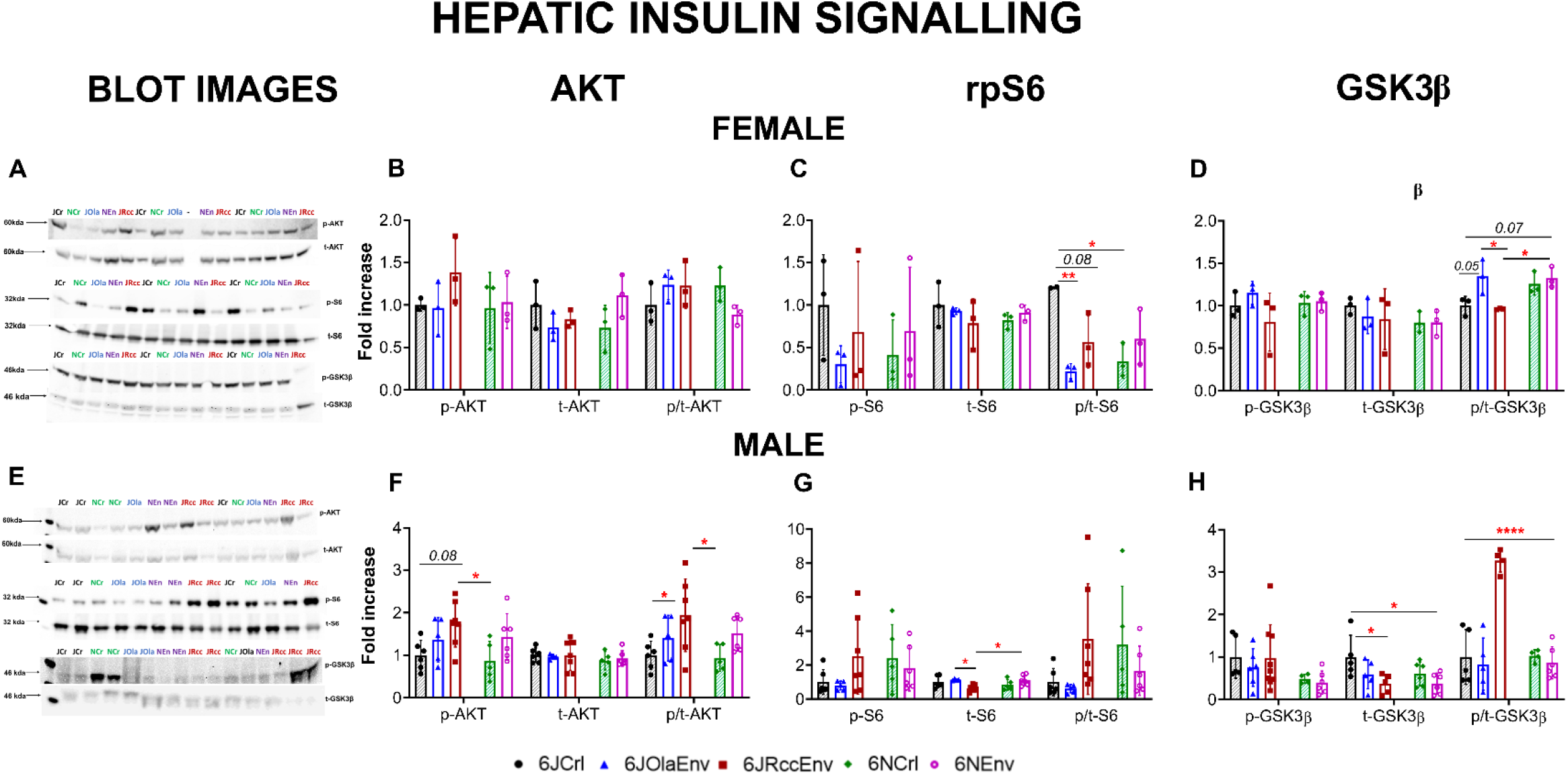
Hepatic insulin signalling markers in C57 substrains of both sexes: Representative images of western blots probed with antibody p-AKT, t-AKT, p-rps6, t-rpS6, p-GSK3β, t-GSK3β for the detection of phosphorylated (p-) or total (t-) AKT, rpS6 and GSK3β in liver of female (**A**) and male (**D**) C57 substrains with molecular weight in kDA. Fold increase in protein levels of p-, t-, p/t ratio of : AKT in female (**B**) and male (**F**); rpS6 in female (**C**) and male (**G**); GSK3β in female (**D**) and male (**H**) C57 substrains, relative to 6JCrl group within each sex displayed. Differences in basal fasted (5h fast as above) hepatic insulin signalling marker levels between substrains within each sex interrogated using one-way ANOVA with Tukey’s post-hoc tests such that 6JRccEnv males had highest levels of p/t AKT and GSK3β while all other substrains apart from 6JCrl females had low levels of p/t-rpS6; sex differences in levels of p/t GSK3β in 6JRccEnv was also noted. All data sets visualised as scatter plots with mean ± SD with significances for displayed on graph as: ****=p<0.0001 vs all other groups, **=p<0.01, *=p<0.05 (Tukey’s post-hoc test), p<0.09 in italics for marginally significant differences. Key for each substrain displayed at the bottom of the figure such that 6JCrl (black), 6JOlaEnv (blue), 6JRccEnv (maroon), 6NCrl (green), 6NEnv (purple) with Crl (Charles River Laboratories) animals of J and N substrain having shaded pattern inside to distinguish from Env (Envigo).

**Fig. 4:**
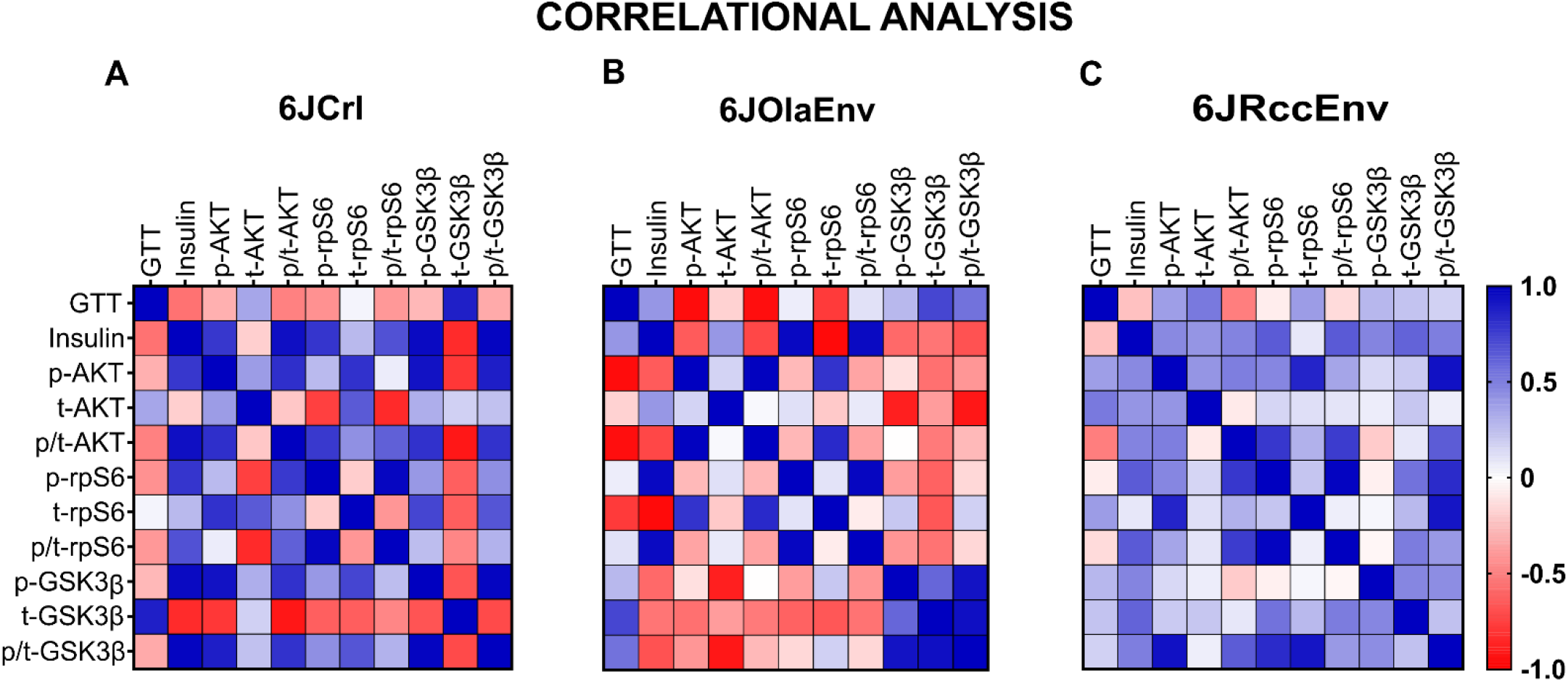
Correlation matrices between GTT glycemic excursion, serum insulin levels and insulin signalling markers in male 6J substrains: Heat plots depict Z-score converted (overall mean) correlation matrices with positive (blue) and negative (red) correlations based on Pearson’s r correlational coefficient, for male 6JCrl (**A**) 6JOlaEnv (**B**), 6JRccEnv (**C**) animals. Upper and lower triangle represent mirror images with GTT indicating area under curve or total glycemic excursion during GTT, insulin indicating basal fasted serum insulin levels followed by hepatic insulin signalling markers as mentioned above. For further detail, see Results.

In male animals, levels of p-AKT in the 6JRccEnv strain were significantly higher than 6NCrl (p=0.04) and marginally higher (p=0.08) than 6JCrl mice. No differences in the levels of t-AKT were observed (F<1). Mirroring the pattern observed in p-AKT levels, 6JRccEnv animals had higher p/t-AKT levels compared to both 6JCrl (p=0.03) and 6NCrl (p=0.03).

Differences were also observed in the levels of p/t-rpS6 in female animals (Fig 3C), such that 6JCrl mice had higher levels compared to most other sub-strains (vs 6NCrl, p=0.01; vs 6JOlaEnv, p=0.009; vs 6JRccEnv, p=0.08). No significant differences were observed in either p-rpS6 (F (4,10) = 0.612, p=0.66) or t-rpS6 (F (4,10) = 0.653, p=0.63).

Unlike female animals, males (Fig. 3G) displayed differences in the levels of t-rpS6. Here, 6JRccEnv had lower levels compared to 6JOlaEnv (p=0.01) and 6NCrl (p=0.03). No changes were observed in the levels of either p-rpS6 (F (4.24) =1.5, p=0.23) or p/t-rpS6 (Welch’s ANOVA, W (4,10.40) = 2.53, p=0.10), although there was a high variability for both markers. While the levels of p-GSK3β (F (4,10) =1.25, p=0.34) and t-GSK3β (F (4,10) = 0.46, p=0.76) did not change in female animals, differences were observed between substrains when examining fold change in p/t-GSK3β (Fig. 3D). Here, 6JOlaEnv had higher levels compared to the other 6J substrains (vs 6JRccEnv, p=0.03; vs 6JCrl, p=0.05). Meanwhile, 6NEnv animals had higher levels of p/t-GSK3β compared to 6JRccEnv (p=0.04) and 6JCr (p=0.07).

In the male cohort (Fig. 3H), no differences between substrains were observed for p-GSK3β (F (4, 27) = 0.94, p=0.45), but 6JRccEnv (p=0.04) and 6NEnv (p=0.03) animals had lower levels of t-GSK3β compared to 6JCrl males. 6JRccEnv had much higher levels of p/t GSK3β compared to all other sub-strains (vs 6JCrl, 6NCrl, 6JOlaEnv, 6NEnv, p<0.0001). A pre-planned comparison between male and female 6JRccEnv for the p/t-GSK3β confirmed that levels were much higher in males (p=0.0004).

### Correlational analysis

To interrogate potential connections between total glycaemic excursion during GTTs (i.e. AUC), with serum insulin levels and molecular markers involved in hepatic insulin signalling (following the format in [22]), we conducted a multi-factorial correlation analysis in 6J male animals. Correlations for 6JCrl, 6JOlaEnv and 6JRccEnv are presented in the main manuscript (see Supplementary Fig. S2 for correlations in 6NCrl and 6NEnv males). Of particular interest is the emergence of clusters comparing different insulin signalling markers with (i) AUC during GTT (ii) serum insulin (iii) between themselves and varying correlation patterns between sub-strains. Our intention was to identify associated potential clusters of interest, rather than searching for individual significances. Individual significances, as well as plots of r and p-values are provided in the supplementary section (Figure S1).

An interesting pattern emerged when comparing GTT AUC values with other markers of interest. Here, a moderate *negative* correlative cluster (red) is observed between AUC and insulin as well as most other hepatic insulin signalling pathway markers only in the 6JCrl animals. The modest *negative* correlation between insulin and AUC in the 6JCrl male animals, although not significant, is interesting as these animals were found to have high glycaemic excursion with low serum insulin levels. Conversely, in the 6JRccEnv animals a weaker, *positive* (blue) cluster was observed instead. The 6JOlaEnv animals meanwhile had an ‘intermediate’ phenotype such that AUC had positive correlations with insulin levels and GSK3β markers but strong negative correlations with other hepatic insulin signalling markers.

Serum insulin levels were found to be highly *positively* (and significantly) correlated to most hepatic insulin signalling markers in the 6JCrl animals and to a *much lesser* extent for the 6JRccEnv animals. Insulin levels in the 6JOlaEnv animals had negative correlations with all hepatic signalling markers besides rpS6. This observation makes sense as male 6JOlaEnv animals had both low serum insulin and rpS6 levels but comparatively higher levels of AKT and GSK3β pathway markers. For serum insulin, correlations between different insulin signalling markers like AKT, rpS6 and GSK3β displayed much stronger positive clusters for the 6JCrl animals compared to either 6JRccEnv or 6JOlaEnv mice, where a loss of these correlations was observed. It was also interesting to observe that levels of t-GSK3β correlated *negatively* with serum insulin and other insulin signalling markers but very *positive*ly with AUC levels (r=0.87) especially in the 6JCrl animals (and to a lesser extent in the 6JOlaEnv cohort). This is also reflected in the male 6JCrl metabolic profile which showed high levels of t-GSK3β with low levels of insulin and other markers (AKT, GSK3β) as well as high AUC during GTT.

## Discussion

Our findings demonstrate that the metabolic profile and glucose clearance of different C57/BL6 substrains are dependent on sex, substrain and supplier, and thus not exclusively influenced by the presence (or absence) of the *Nnt* mutation. 6JCrl and 6NEnv of both sexes and male 6JRccEnv mice displayed the highest impairments in glycaemic excursion compared to other substrains. Indeed, glycaemic excursion was modulated not just by substrain but also supplier and sex. For animals supplied by Charles River, 6J was marginally more affected compared to 6N with no differences between sexes. This was not the case for Envigo animals where the glucose intolerance profiles of the substrains differed between sexes (6NEnv>6JRccEnv/6JOlaEnv: females; 6NEnv/6JRccEnv>6OlaEnv: males), sometimes even within the same substrain (6JRccEnv males > 6JRccEnv females). In male Envigo animals, despite 100% genetic concordance between the 6JRccEnv and 6JOlaEnv animals as per Envigo’s technical data sheet, the former was found to be more glucose intolerant compared to the latter [19,26,27].

Differences between substrains, sexes and vendors may explain why previous publications have reported inconsistent results with regards to GTT results/data of 6J and 6N animals, further complicated by the omission of sex reporting in some studies [11–13,28].Only a few have compared GTT responses on a standard chow diet, as was the case here. It is also interesting to note that the 6N substrains were not found to be different from each other, irrespective of vendor or sex, confirming previous results [19].

The present study also provides important information regarding fasted serum insulin levels. Prior publications have indicated that although 6J (Jackson Laboratory, sex missing) have impaired glucose tolerance, they have normal insulin sensitivity. These mice, however, exhibited decreased insulin secretion, including a resistance to elevations in circulating insulin after a short-term HFD [5,11]. Accordingly, we did not detect elevations in serum insulin levels in 6JCrl mice of both sexes despite the presence of glucose intolerance, while male 6JRccEnv mice displayed high fasted insulin levels, which could explain their glucose intolerance. The two distinct metabolic phenotypes could explain impairments in GTTs – one, where low basal insulin levels results in impaired glucose clearance during a GTT (as seen in 6JCrl; both sexes) and the other, where sustained elevations in fasted basal insulin levels leads to glucose intolerance (as in male 6JRccEnv animals) [29].

Previous research implicated an *Nnt* mutation in the 6JCrl mice in their glucose intolerance and impaired insulin secretion. It is clear from the results presented here that *Nnt* deletion is not the only factor responsible for metabolic alterations as substrains with intact *Nnt* (Table 1) also showed glucose intolerance or high insulin levels. Therefore, the *Nnt* mutation at best moderately contributes to disturbed glucose handling [4,14,26]. Instead, basal fasted hepatic insulin signalling differed between substrains in both sexes, a likely contributor to variations in metabolic phenotypes observed. For example, p/t-AKT ratio was found to be greater in male 6JRccEnv animals cf other substrains obtained from Charles River (6J/6N). Increased liver phosphorylation of AKT has previously been associated with higher fasted plasma insulin levels and hyperinsulinemia in C57BL6 animals on a HFD, although no information about substrain, vendor or sex was provided [30]. Like AKT, an increase in p/t-GSK3β levels was also noted in the 6JRccEnv males, where a difference between sexes was also noted. Incidentally, sex differences in hepatic ischemia/reperfusion injury have been found to be mediated by a male specific gene SRY (sex determining region on the Y chromosome) through the upregulation of GSK3β phosphorylation [31]. Increased GSK3β activation has been linked to glucose intolerance and insulin resistance, which may explain phenotypes observed in male 6JRccEnv animals [32–34].

In case of rpS6, substrain differences in p/t-rpS6 were observed only in females, and 6JCrl animals presented with the highest ratio. This sex-specific increase in phosphorylation, possibly driven by increased S6K1 activity, could indicate a compensatory mechanism in response to low fasting insulin levels and impaired glucose clearance to initiate further protein translation. In fact, S6K1 deficient mice have been found to be hypo-insulinemic and glucose intolerant, yet are insulin-sensitive and protected against HFD-induced obesity and hepatic steatosis [35–37].

Finally, results from the multi-factorial correlational analysis in male 6JCrl, 6JOlaEnv and 6JRccEnv animals provide further evidence of substantial differences in metabolic profiles. Here, 6JCrl animals exhibited much stronger and even opposing correlations (for e.g., GTT-AUC) compared to 6JRccEnv animals for glucose homeostasis, fasting insulin levels and insulin signalling markers, with 6JOlaEnv animals exhibiting an ‘intermediate’ phenotype which had some similarities to both 6JCrl and 6JRccEnv.

Taken together, 6J and 6N substrains from the same vendor were found to exhibit different metabolic profiles, in accordance with previous literature. These differences were stronger for animals from Envigo compared to Charles River, as we report sex differences even within a particular substrain. The profile of 6JCrl mice of both sexes may be better suited for metabolic studies as they are glucose intolerant, with low fasting insulin levels. In comparison, the 6N substrains tested do not exhibit considerable difference between them or between sexes, irrespective of vendors. Specifically, 6NCrl animals of either sex did not present with basal metabolic phenotypes, which would make them a good choice as healthy controls for a variety of biomedical experiments unless impacted by the retinal degradation gene.

6JRccEnv mice, especially males, presented with glucose intolerance, elevated plasma insulin levels and phosphorylation of insulin signalling markers. Therefore, diabetes or obesity research designed using these mice must bear in mind that their basal metabolic profile is different from other C57/BL6 substrains. The large increase in basal fasted phosphorylation of GSK3β could also have important implications for dementia research since GSK3 overactivity is implicated in the pathogenesis of tau [38].

Finally, our results call for great caution and sufficient inclusion of control experiments for any research where metabolic function may contribute to outcomes. A full characterisation is needed to understand implications on long-term interventions or creation of new transgenic animals. Additionally, sex differences in metabolic profiles indicate the need to consider sex as a biological variable more strongly in metabolic research, as a moat studies use only male animals [39]. In line with recent initiatives [1,20,21] we strongly recommend that researchers fully and transparently report the sex, substrain and supplier of experimental animals, and consider the use of littermates raised under matching conditions to improve credibility and reproducibility of biomedical research.

## Supporting information

Supplemental figures 1 and 2 with legends

## ACKNOWLEDGEMENTS

We would like to acknowledge the staff of the Medical Research Facility for their support with animal care, handling and GTT experiments and members of the Riedel/Platt lab for their assistance with tissue collection.

