## Supplemental figures 1 and 2 with legends for "How stra(i)nge are your controls? A comparative analysis of metabolic phenotypes in commonly used C57 substrains"

**SUPPLEMENTARY INFORMATION**

**Fig. S1:** **Correlation matrices between GTT glycemic excursion, serum insulin levels and insulin signalling markers with values of correlational coefficients (r) and p-values indicated:** Heat plots depict Z-score converted (overall mean) correlation matrices with values of the Pearson’s correlational coefficients (positive (blue) and negative (red)) indicated within each of the matrices on top for 6JCrl (**A**) 6JOlaEnv (**B**) 6JRccEnv (**C**) male animals and p-values on the bottom (<0.09) for 6JCrl (**D**) 6JOlaEnv (**E**) 6JRccEnv (**F**), respectively. Upper and lower triangle represent mirror images with GTT indicating area under curve or total glycemic excursion during GTT, insulin indicating basal fasted serum insulin levels followed by hepatic insulin signalling markers as mentioned above. For further detail, see Results.

**Fig. S2: Correlation matrices between GTT glycemic excursion, serum insulin levels and insulin signalling markers in male 6N substrains**: Heat plots depict Z-score converted (overall mean) correlation matrices with positive (blue) and negative (red) correlations based on Pearson’s r correlational coefficient for male 6NCrl (**A**) and 6NEnv (**B**) animals. Upper and lower triangle represent mirror images with GTT indicating area under curve or total glycemic excursion during GTT, insulin indicating basal fasted serum insulin levels followed by hepatic insulin signalling markers as mentioned above. For further detail, see Results.

**Figure S1**

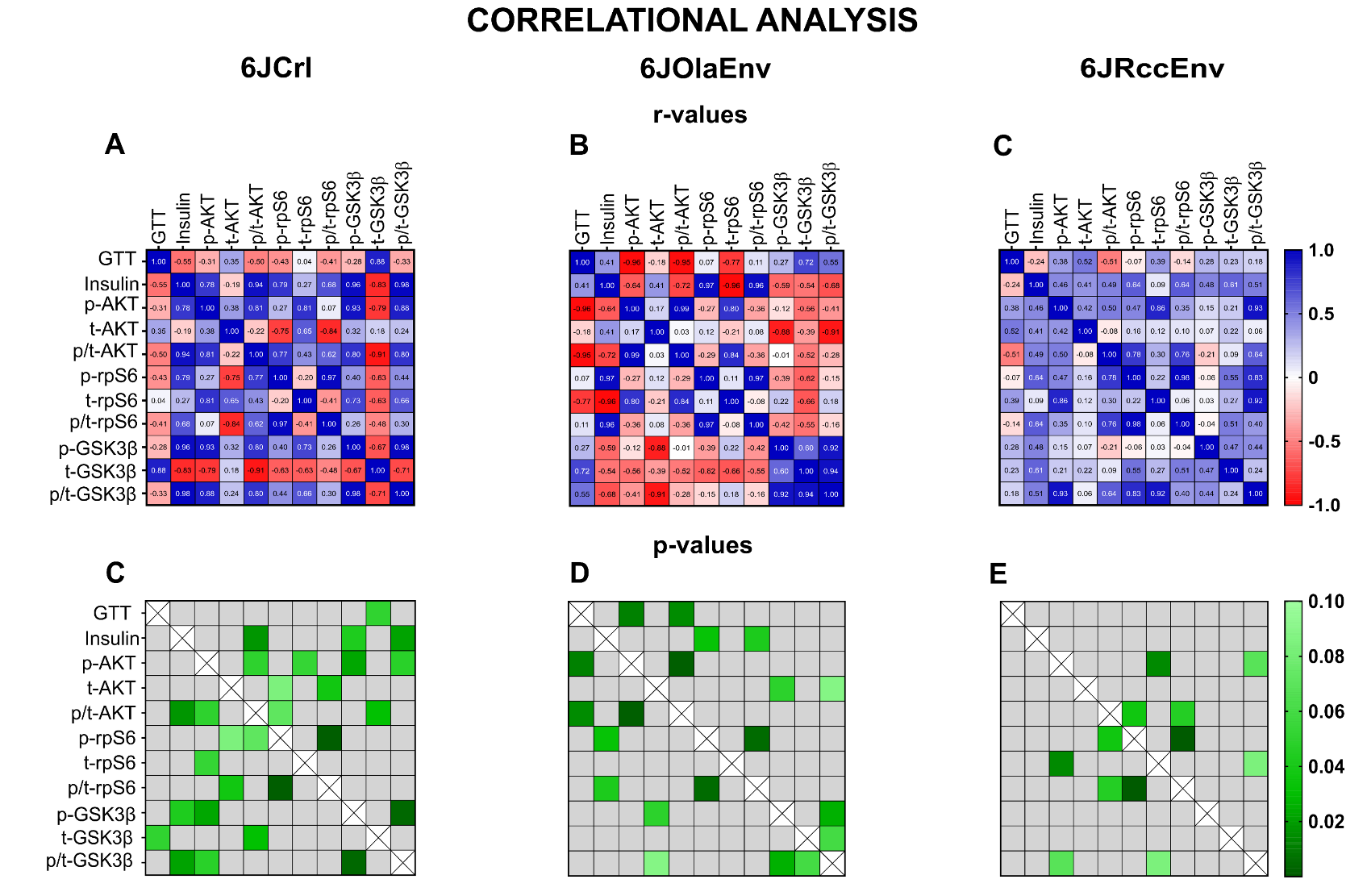

**Figure S2**

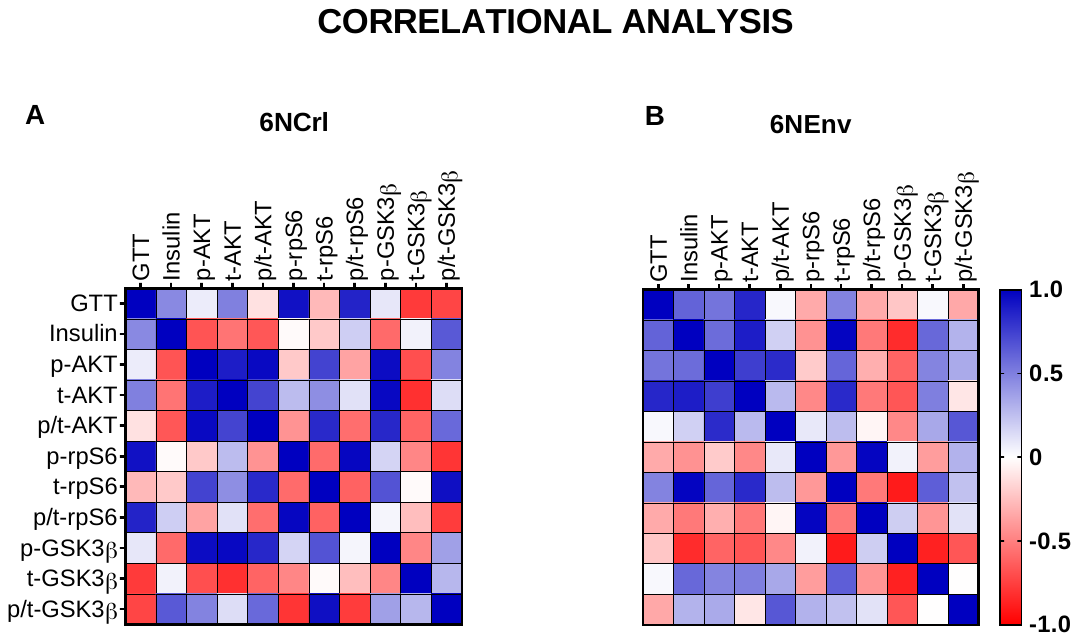
